# Adaptive oscillators provide a hard-coded Bayesian mechanism for rhythmic inference

**DOI:** 10.1101/2022.06.18.496664

**Authors:** Keith B. Doelling, Luc H. Arnal, M. Florencia Assaneo

## Abstract

Bayesian theories of perception suggest that the human brain internalizes a model of environmental patterns to reduce sensory noise and improve stimulus processing. The internalization of external regularities is particularly manifest in the time domain: humans excel at predictively synchronizing their behavior with external rhythms, as in dance or music performance. The neural processes underlying rhythmic inferences are debated: whether predictive perception relies on high-level generative models or whether it can readily be implemented locally by hard-coded intrinsic oscillators synchronizing to rhythmic input remains unclear. Here, we propose that these seemingly antagonistic accounts can be conceptually reconciled. In this view, neural oscillators may constitute hard-coded physiological priors – in a Bayesian sense – that reduce temporal uncertainty and facilitate the predictive processing of noisy rhythms. To test this, we asked human participants to track pseudo-rhythmic tone sequences and assess whether the final tone was early or late. Using a Bayesian model, we account for various aspects of participants’ performance and demonstrate that the classical distinction between absolute and relative mechanisms can be unified under this framework. Next, using a dynamical systems perspective, we successfully model this behavior using an adaptive frequency oscillator which adjusts its spontaneous frequency based on the rate of stimuli. This model better reflects human behavior than a canonical nonlinear oscillator and a predictive ramping model, both widely used for temporal estimation and prediction. Our findings suggest that an oscillator may be considered useful as a potential heuristic for a rhythmic prior in the Bayesian sense. Together, the results show that adaptive oscillators provide an elegant and biologically plausible means to subserve (bayesian) rhythmic inference, thereby reconciling numerous empirical observations and a priori incompatible frameworks for temporal inferential processes.

## Introduction

According to Bayesian accounts of perception, the brain internalizes and deploys a predictive model of the environment, based on prior experience. For example, in adverse conditions, the context of a sentence can either bias word recognition^1^ or otherwise enhance the perception of the words that would be incomprehensible out of context^2^. Such predictive models proactively guide perception^3–5^, actions^6^, and decision-making^7^ by biasing internal representations towards top-down expectations and thereby reducing internal variance. In these *forward* models, predictions essentially arise from higher-level hierarchical brain regions that propagate downward to constrain sensory processing of incoming events in sensory areas. As such, forward models are often considered as reflecting individuals’ prior sensory experience over their lifetime. Whether intrinsic neurobiological features (e.g. oscillations) pre-equip internal models with hard-coded priors to constrain perceptual analysis of upcoming events is generally overlooked.

Tracking the rhythm of a sequence of events – as in speech or music – reduces temporal uncertainty and facilitates the processing of upcoming events^8^. Such temporal predictions could clearly benefit from a similar Bayesian framework. However, most empirical works in this domain have focused on the potential role of neural oscillations as a neurophysiological substrate for predictions in the time domain^9–12^. In this view, neural oscillators synchronize their excitability phase with external sequences, thereby reducing internal noise and optimizing the processing of incoming events^13–15^. In this sense, the phase of a neural oscillation can be used as an index for prediction in time, a mechanism that may be considered as constitutive of the inferential process. Although readily applied in the context of isolated intervals^16^, Bayesian perceptual accounts are seldom considered to account for anticipatory processing of *sequences* of events in the time domain (although see this recent proposal^17^).

Here, we bring together these parallel lines of research to pose the following question: Can neural oscillations and Bayesian accounts be adjudicated between or must they be reconciled when it comes to rhythmic sequence processing? While Bayesian models are often considered in the sense of long-range top-down or lateral information flow, they could also be supported by the more local dynamics afforded by neural oscillators, essentially hard-coding prior expectations for rhythmicity in sequences. In this view, hard-coded physiological mechanisms (e.g. neural oscillations) might reflect intrinsic neurophysiological adaptations to the statistics of the environment.

One issue that arises in adjudicating between underlying models in rhythm perception is that all reasonable models will yield the same prediction under *perfect rhythmicity*. However, natural stimuli generally possess regular temporal statistics often construed as pseudo-rhythmicity: ecological sounds are rarely *perfectly* rhythmic and instead possess wide variability in timing^18–20^. Using temporally variable sequences is arguably necessary to capture the essence of internal models, namely the extraction of noisy patterns to reduce uncertainty about future events. As such, by introducing temporal uncertainty – i.e. jittering the events – in the sequence, we can better distinguish between models: variance in the intervals leads to variance of predictions in the models.

Variance in timing also arguably represents a challenge for the neural oscillatory synchronization hypothesis^21^. Such mechanisms could become less efficient (or even detrimental) in the context of more irregular temporal patterns. Recently, we theoretically demonstrated that even in the face of such variability a basic oscillator can still synchronize to a variable rhythm^22^. Does this kind of synchronization still work as a mechanism for human temporal prediction without perfect isochronicity? Can the phase adjustments that lead to successful synchronization be reconciled with a Bayesian account of perception? To answer these questions, we devised temporal sequences with regular statistics but with clear temporal jitter. Participants therefore could predict when the next tone was most likely to occur without being able to have total confidence. This protocol allowed us to determine how temporal expectations are executed under these adverse - but more naturalistic - conditions.

While previous studies have used similar experimental designs, the algorithm driving the subject’s responses was taken for granted and the analyses were conducted according to this assumption despite the fact that other algorithms may yield different results. In this line, two main distinct algorithms have been proposed in the literature to subserve perceptual timing: absolute or relative timing^23^. Absolute timing relies on the estimation of the concrete duration of discrete time events. Relative timing, instead, refers to the computation of a time interval with respect to a standard defined by a temporal regularity present in the environment. Typically, it is assumed that one or the other time assessment drives behavior according to the specificities of the experimental design. For example, it has been hypothesized that timing of intervals in irregular time sequences recruits absolute timing mechanisms while regular sequences prefer relative timing ones^24^. Here, in contrast with previous studies, we do not presuppose that one or another timing mechanism will drive participants’ expectations. As an alternative, we adopt a data driven approach, where the timing algorithm is determined by the participants’ responses. By taking this feature into account, we develop a more complete understanding of human behavior and supplement this understanding with models at the computational and algorithmic levels to better understand how listeners behave and whether neural oscillatory models can still reflect human behavior in this more naturalistic condition.

Our findings show that what seems to be two different timing mechanisms can be different sides of the same coin. While participants’ responses reflected different algorithms subserving perceptual timing depending on the temporal features of the acoustic stimulus, we show that the whole pattern of responses can be captured by a single mechanism. First, we show that participants’ behavior is consistent with Bayes’ Theory with regards to temporal prediction. Then, we introduce a biophysical neural model capable of explaining participants’ responses. More precisely, we propose an adaptive frequency oscillator as a reasonable candidate for temporal prediction under these more ambiguous circumstances by taking qualities typically used in other models of the literature (e.g. predictive ramp) to improve performance of the oscillator model. Taken together, we show the advantage for proponents of oscillatory models to develop more complexity in their theories to accommodate realistic scenarios of human perception. At the same time, our findings support the use of oscillatory components as a hard-coded rhythmic prior in the Bayesian sense.

## Methods

### Participants

In total, four behavioral experiments were collected running the same task design except for the average tone rate (1.2 Hz, 1.2 Hz – low jitter, 2 Hz and 4 Hz). Across all experiments, we collected data from 215 participants (1.2: 35, 1.2LJ: 37, 2: 78, 4: 65) in a mixture of in person and online experiments (1.2: online, 1.2LJ: online, 2: in person, 4: in person; cf. COVID). Of these, 44 participants were removed due to performance not significantly above chance making for a total of 171 (101 females; mean age, 23 years; age range, 19 to 36 years) participants collected in the study (1.2: 34, 1.2LJ: 33, 2: 58, 4: 46). Participants were given course credit for their participation at NYU. The study was approved by the University Committee on Activities Involving Human Subjects (UCAIHS) at New York University.

### Experimental Design

Participants listened to a series of tones and were asked to identify if the last tone was earlier or later than expected. Participant responses were given by 4 possible button responses: two buttons to indicate confident early or late responses and two in between to indicate less confidence. For our analysis, we group all responses regardless of confidence.

On each trial, participants hear *N* tones followed by a final probe tone about which participants give their responses, where *N* is randomly chosen on each trial between 8, 9 or 10 (see Figure 1a). A within-trial average period between tones is selected to define the expected location of each tone. This event period *T* is drawn from a uniform distribution on each trial whose bounds ([*b_l_*, *b_h_*]) were set for each experiment (1.2 Hz: [700, 1000] ms, 1.2 Hz low jitter: [700, 1000] ms, 2 Hz: [400, 600] ms, 4 Hz: [210, 290] ms). Each tone (*t_i_*) is then displaced from the expected location by a gaussian random variable *ε* ~ *Norm*(0, σ) where σ is approximately 20% of the mean period across the experiment (1.2 Hz: 170 ms, 2 Hz: 100 ms, 4 Hz: 45 ms), with the exception of the low jitter version of the 1.2 Hz experiment which sought to test the effect of a reduced temporal jitter (1.2 Hz low jitter: 45 ms). In all cases, the final probe tone (*t_probe_*) is placed at the following expected location where the temporal error is drawn from a uniform distribution.

**Figure 1.**
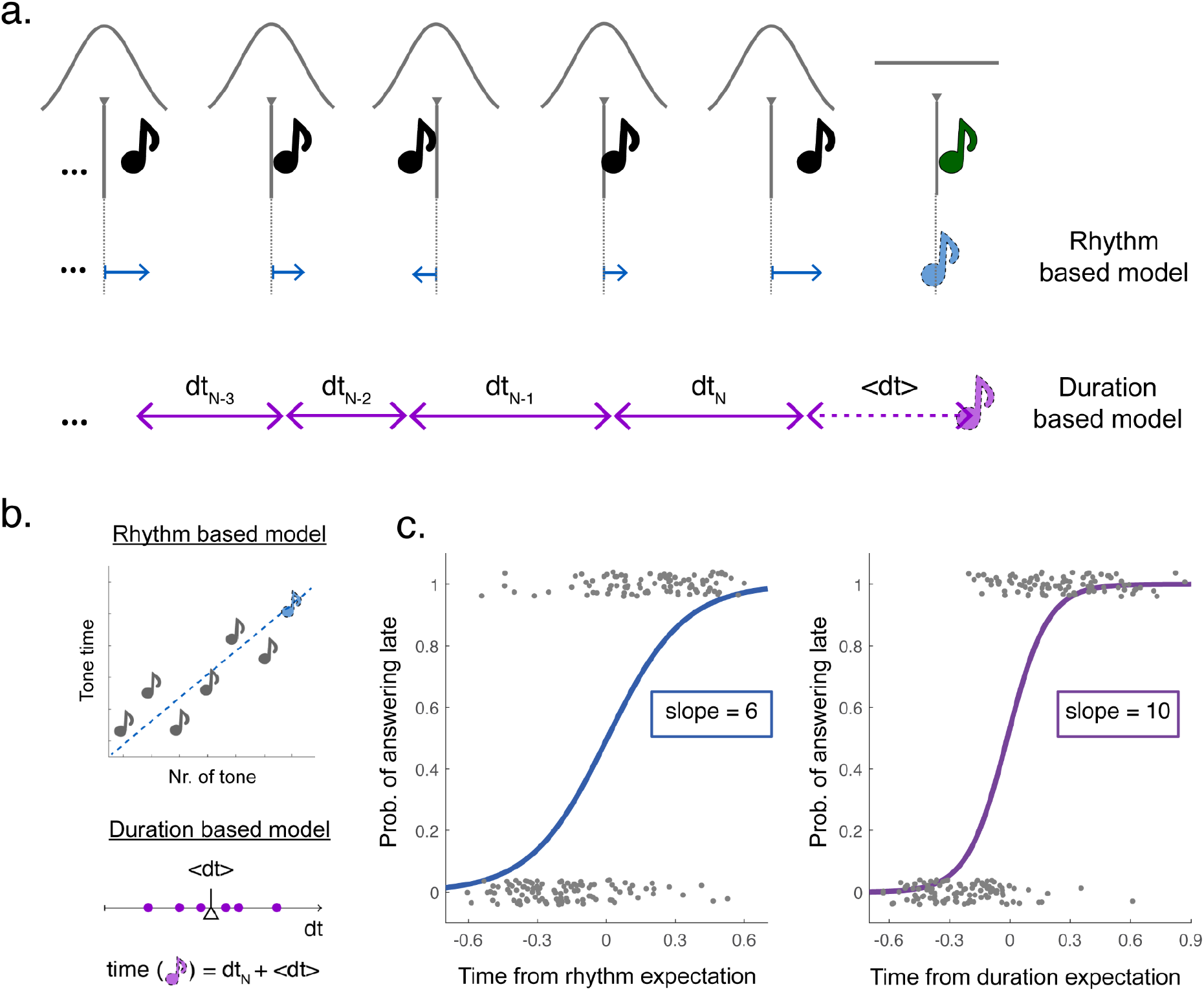
Experimental design and analysis framework. a) Schema reflecting the task design and the different algorithms used to estimate the expected final location. Each cue tone (black) is drawn from a gaussian distribution (gray line) with expected mean in with rhythmic placement. The final probe tone (green) is drawn from a uniform distribution around the next expected time. The rhythm (blue) and duration (purple) based algorithms minimizes the error from expected locations or averages the collected time delays between tones respectively. b) Computations used to calculate rhythm and duration algorithms. Rhythm uses linear regression, fitting a line to predict timing of the next tone based on its position in the sequence. Duration stores the intervals between tones, averages and adds this interval to the final tone location. c) Analysis of example participants. Responses (coded 0 for early, 1 for late) are compared to both the rhythm algorithm (left) and the duration algorithm (right). Logistic model is fitted to the results and the slope of the fit is used as a measure of consistency with the algorithm. This example shows a participant more consistent with the duration method.

Mathematically, the parameters defining each trial can be defined by the following set of equations, where *U* represents a uniform distribution, *Norm* a gaussian one and *E* the expectation of a random variable:

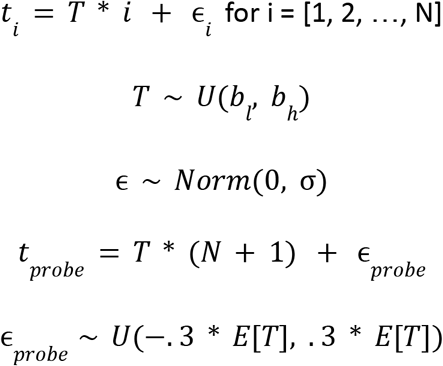

Each experiment comprised 150 trials and each participant was assigned to one of the 4 experiments: 1.2 Hz low jitter, 1.2 Hz, 2 Hz or 4 Hz.

### Experimental Analysis

#### i. Duration vs Rhythm

While the tones are generated by a specific set of equations, the listener does not have access to this generating algorithm and must therefore infer it on the basis of very limited data (10 tones at most). In fact, the task can be solved by different algorithms leading to different results as to whether the final tone is early or late. We considered two possible algorithms in particular that: 1) seem the simplest possible choice, 2) represent two different strategic goals and 3) align with the two typically proposed timing mechanisms (i.e., absolute and relative timing). They are exemplified in **Figure 1ab.** The *Rhythm* algorithm assumes that participants use the sensory evidence only to infer where the expected (mean) location would have been, while the *Duration* algorithm takes sensory evidence at face value and adds the mean interval on to the timing of the previous tone. The main difference between these algorithms is whether you should allow for drift in the expected locations. In that sense, the Rhythm algorithm resembles *relative timing* given that the time of the incoming tone is predicted by computing the overall temporal regularity in the preceding sequence. In contrast, the Duration algorithm compares to *absolute timing*, since it is the absolute time that goes by between tones that drives prediction.

A primary question for our study was to uncover, given a tone sequence, how do participants navigate this distinction and deal with the lack of a ground truth to generate a response. To answer this, we score participant data by *both* possible mechanisms to help us identify which one the participants behavior most closely resembles. Probe tones are coded by their deviation from either the Duration prediction or the Rhythm prediction. Then, we fit two sigmoid functions between this deviation (from the two algorithms) and the response of the participant as to whether the tone is early (recorded as 0) or late (1) as in **Figure 1c**. We then extract the slope parameter of this fit as a measure of the precision of their performance for either algorithm and compare the two obtained slopes to see which algorithm performs best in terms of sorting the participants’ responses. Figure 1c shows an example of a subject who performs best relative to the duration algorithm. For analyses merging the data of different experiments (i.e., different rates) the normalized logistic regression were estimated. Meaning that the time difference between the probe tones and the Duration and Rhythm predictions were divided by the mean SOA of the corresponding trial.

Logistic regressions on each participant’s responses were fitted using the fitglm matlab function, which returns a generalized linear model fit. Participants with a noisy response pattern (χ^2^ statistics against constant model, p>0.05) were excluded from the follow up analyzes. The number of participants excluded for each condition are: 18 from a total of 65 for the 4 Hz condition, 24 from a total of 78 for the 2 Hz and 2 from a total of 72 for the 1 Hz condition (including both low and high jitter).

#### ii. Statistics

All between- and within-subject comparisons were assessed by nonparametric Mann-Whitney-Wilcoxon and Wilcoxon signed-rank tests, respectively.

The relationship between the mean of the slopes for Rhythm and Duration and their difference was investigated and the relationship fit by a 2nd-order polynomial to estimate how overall performance relates to method behavior. The model is fitted using a least-squares estimation using the matlab polyfit function. Confidence Intervals of the parameter estimates were assessed using the polyconf function to assess the statistical significance of the parameters relative to 0. And the overall goodness of the fit was estimated by means of the adjusted R-square, penalizing the variables addition to the model.

### Computational modeling

We designed two model types to explain the behavioral effect in terms of computational processes and internal dynamics that could generate them. First, we designed a Bayesian model to understand how the combination of expectation and sensory measurement might yield similar human behavior. Next, we designed an adaptive frequency oscillator as a candidate neural mechanism to underlie this mechanism.

### Bayesian Model

Our Bayesian model makes several assumptions. First, estimates of the timing of an event are measured with gaussian noise, such that if the i-th tone occurs at time *t_i_*, then the model’s representation of this time will be drawn from a gaussian distribution *N*(*t_i_*, σ*_s_*) where σ_*s*_, the standard deviation of the gaussian, represents sensory noise or inversely the precision of the model representation. Next, we assume that this noisy estimate of time is combined with a prior expectation of timing on the basis of preceding tones. We devise this prior in two possible methods following the same Duration-based and Rhythm-based models we used in the behavioral analysis. The Rhythm-based prior uses linear regression with unequal variance to estimate the mean and variance of the next expected location. Thus, the mean expected timing of the next tone can be calculated with the following equation assuming t=0 is defined as the start of the first tone.

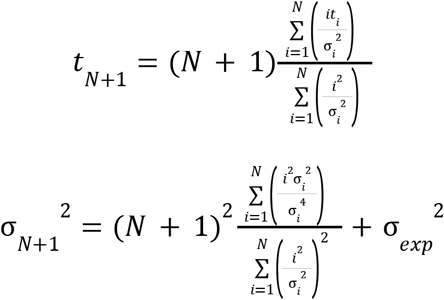

Where *i*, is the index of each tone, *t_i_* is the true time of each tone, σ_*i*_ is the standard deviation of the gaussian representing the observer’s uncertainty and σ_*exp*_ is the uncertainty due to the experiment, the standard deviation of the jitter applied to each tone which we assume the participant knows *a priori*.

The duration-based prior calculates the difference between neighboring tone times to estimate the mean interval between tones and adds this interval to the final tone time to generate an estimate of the prior. The mean and variance are calculated as follows:

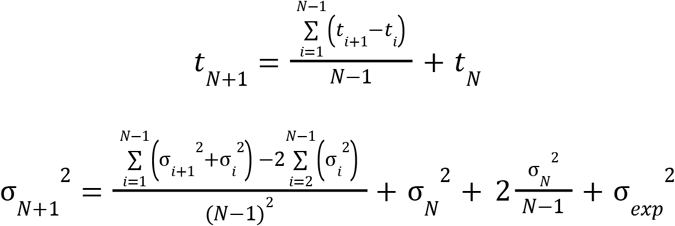

Where the terms 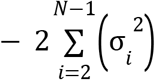 and 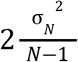 account for the covariance between neighboring intervals and between the mean interval and the final tone time, respectively. The posterior distribution is calculated as a multiplication between the two gaussian distributions of sensory measurement and prior expectation.

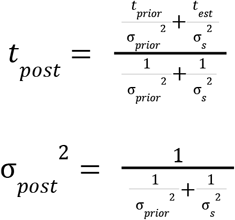

Both priors, the Rhythm-based and the Duration-based, apply only after the two tones have been presented. For the first two tones, we assume a flat prior such that the posterior distribution is equal to the sensory measurement, *N*(*t*|*t_i_*, σ_*s*_).

This model setup yields a posterior distribution for the estimate of each tone within the sequence. Note that the posterior for each tone is based purely on preceding tones and is not updated on the basis of later information.

Finally, the model must make a decision as to whether the final tone is earlier or later than expected. In this case, rather than combining prior expectation and sensory measurement, we compare the two distributions to assess the probability that the sensory estimate is higher than the prior expectation by marginalizing over time the product of cumulative distribution of the prior and probability distribution of the sensory estimate.

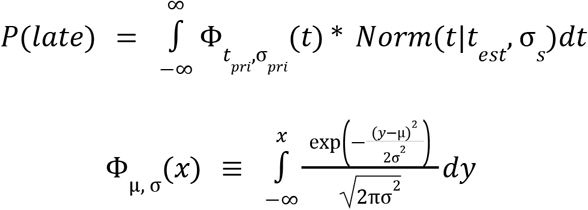

Using this probability, *P*(*late*), we flip a weighted coin on the basis of the probability distribution to determine if the model responds late or early to the trial, and analyze the output exactly as described in the data analysis section on human behavior. We then investigate the model’s behavior by roving the amount of noise in the sensory estimate σ_*s*_ to see how the model behaves with more or less sensory noise.

### Adaptive Frequency Oscillator

The Wilson-Cowan (WC) model is a biophysically inspired neural mass model which has been widely used in the literature and shows a rich set of possible dynamics^25,26^. This model assumes that a given brain region is composed by an excitatory and an inhibitory neural population interacting with each other; and can be described by the following set of equations:

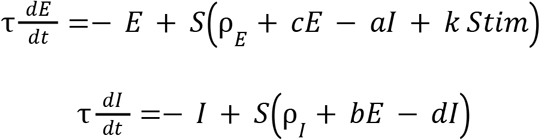

Where *E* represents the excitatory population, *I* the inhibitory one, *S* is a sigmoid function, *a,b,c* and *d* represent the synaptic coupling, τ is the membrane time constant, *Stim* is the external stimulus driving the brain region activity, *k* is the strength of the coupling between the brain and external stimulus, *and* ρ_*E*_ and ρ_*I*_ are stable inputs that the different populations receive from distant brain areas. In the current work, a model like this has been adopted to represent auditory regions and the input to the excitatory population has been assumed to be proportional to the broadband envelope of the auditory stimulus:

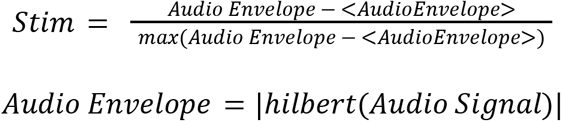

Importantly, the set of equations defining the WC model depict different dynamics for different combinations of (ρ_*E*_, ρ_*I*_). Namely, different bifurcations take place for different trajectories in the (ρ_*E*_, ρ_*I*_) space (for more details see pp. 47 of this detailed analysis^26^). In previous work^22,27^, where a neural oscillator is hypothesized, the parameters (ρ_*E*_, ρ_*I*_) were fixed and set close to an Andronov-Hopf bifurcation where the system behaves as an oscillator with a fixed natural frequency (e.g., see also Figure S1). We refer to this selection of parameters as the classic oscillator. Here instead, in order to get an adaptive frequency oscillator, we assigned the following dynamics to the parameters:

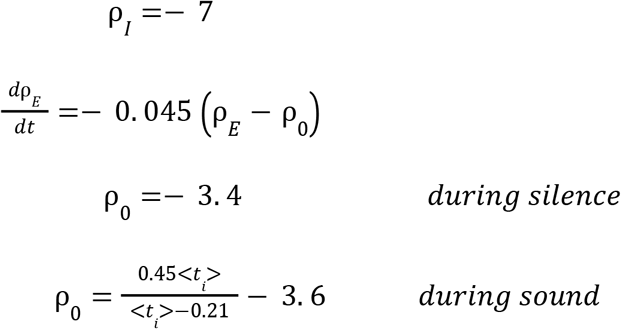

Where *t* represents the period of the perceived sound (i.e., the time interval between sounds is estimated and the mean value is actualized right after each tone). In order to select the functional form for ρ_0_, we numerically explored the relationship between the natural period of the system and ρ_*E*_ (see Figure 2). The functional form for ρ_0_ has been chosen to drive the system to match the perceived auditory period. The chosen dynamics places the system during silence at rest (ρ_*E*_ = − 3.4) and close to a saddle node in limit cycle bifurcation. When the sequence of tones begins ρ_*E*_ evolves in time through ρ_0_, crossing the bifurcation and allowing the system to adjust the natural frequency of the oscillator in order to match the rhythm of the stimulus.

**Figure 2.**
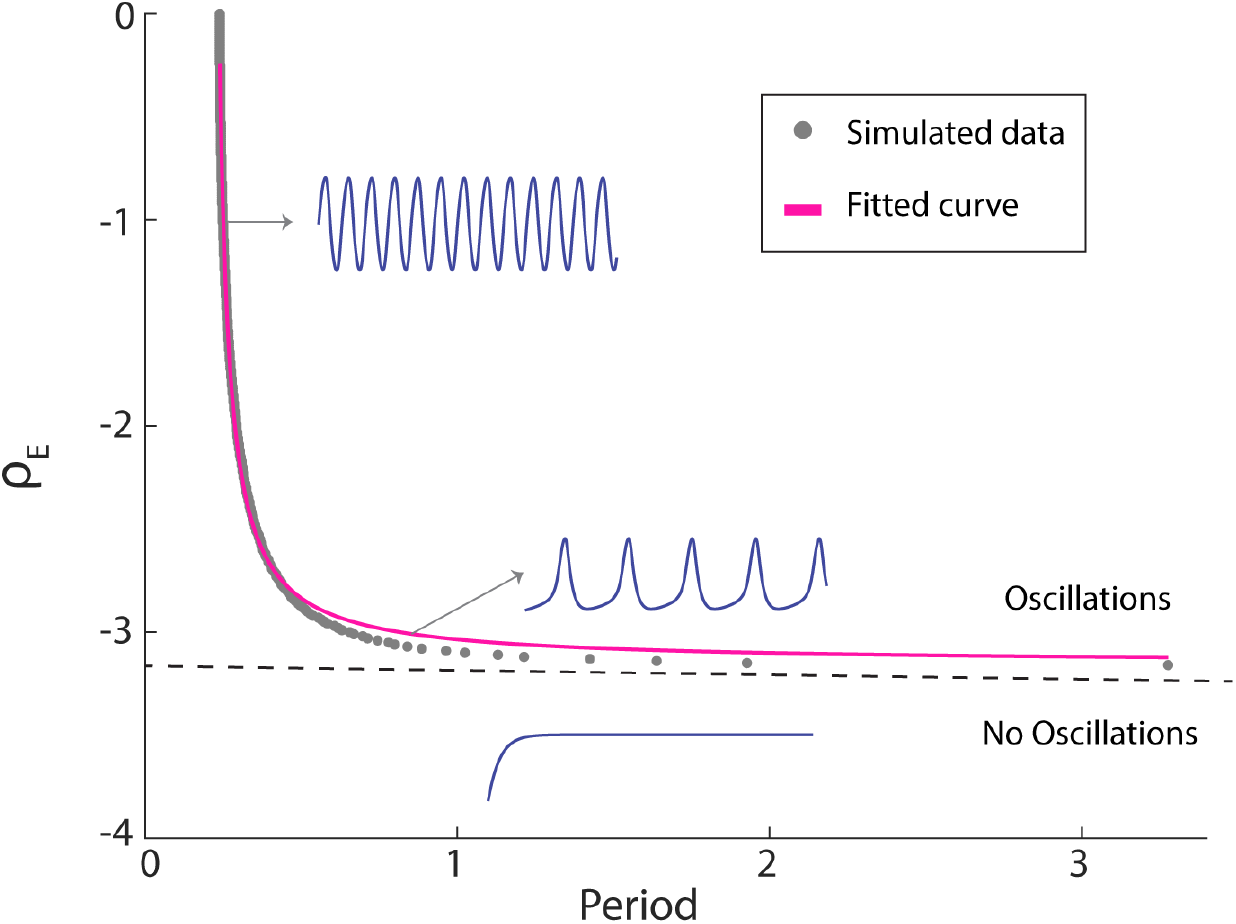
Oscillatory behavior of the WC model. Running numerical simulations of the WC equations with no stimulus (*Stim=0*), we estimated the period of the excitatory population activity for different values of ρ_*E*_ (gray dots). A saddle node in limit cycle bifurcation takes place around ρ_0_ =− 3. 16 (dashed line). Such a bifurcation gives birth to very slow oscillations rapidly increasing its frequency as the relevant parameter departs from the bifurcation point. We fitted a rational function to the numerical data to get an analytic parametrization of ρ_*E*_ as a function of the natural period of the system. Blue traces depict the activity of the excitatory population in the different regimes

### Simulated participant’s answers

To simulate behavior with the AFO the parameters of the model were set to: *a*=*b*=*c*=10, *d=-2* as typically selected in the literature^26^, τ = 1/17 and ρ_*I*_ =— 7. Different participants were modeled by varying the coupling value (*k* = 0. 2 + *U*(− 0.1, 0.1)) and 300 different auditory trials were evaluated per participant.

The time interval between trials was assigned randomly as: *ITI* = 0. 75 + *U*(− 0.25,0.25) and stimuli were generated in the same way as for the behavioral protocol, but removing the probe tone. For each trial the phase of the AFO at the time where the probe should appear was estimated as the phase of the Hilbert transform of the excitatory population activity. The AFO performance was computed by fitting a logistic regression to adjust early or late responses (0 or 1) according to Rhythm and Duration algorithms as a function of the oscillators phase (using (*cos*(θ), *sin*(θ)), for each simulated participant (i.e., for each set of 300 phases computed as stated before). As well as for the experimental data, simulated participants with a noisy response pattern (χ^2^ statistics against constant model, p>0.05) were excluded.

In Fig. S1, in addition to the above simulation, we also use phase concentration to assess what information is contained within the base form of the Wilson-Cowan Model. We use the Hilbert transform to estimate phase of the excitatory population activity at the *expected* time as predicted either by Rhythm or Duration across 300 different trials and use phase concentration to estimate how consistent this phase is across trials. Higher concentration infers that the phase is synchronized to this expected timing. Phase concentration is assessed using the following equation:

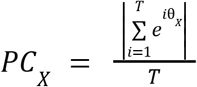

All simulations were run in MATLAB using a standard Euler solver with a time step of 0.0001 sec.

## Results

### Human Behavior

We first compared human responses relative to our two algorithms across the three distinct stimulus rates used in our three experiments. Figure 3a reveals the results of this analysis. Stimulus rate significantly affected algorithm performance. At 1.2 Hz, participants significantly preferred the Duration algorithm, showing higher (steeper) slopes for this compared to Rhythm (Wilcoxon signed-rank test, two-sided p < 0.001). Instead, participants in the 2 Hz and 4Hz study showed a significant preference for the Rhythm algorithm, with higher slopes for Rhythm compared to the Duration algorithm (Wilcoxon signed-rank test, two-sided 2 Hz: p=0.004 and 4 Hz: p<0.001). Furthermore, Slope differences (i.e., Slope_DUR_– Slope_RHY_) significantly differ between studies (Mann-Whitney-Wilcoxon test, two sided, Bonferroni corrected for 3 comparisons: p_1.2Hz-2Hz_<0.001, p_1.2Hz-4Hz_<0.001, p_2Hz-4Hz_= 0.028).

**Figure 3.**
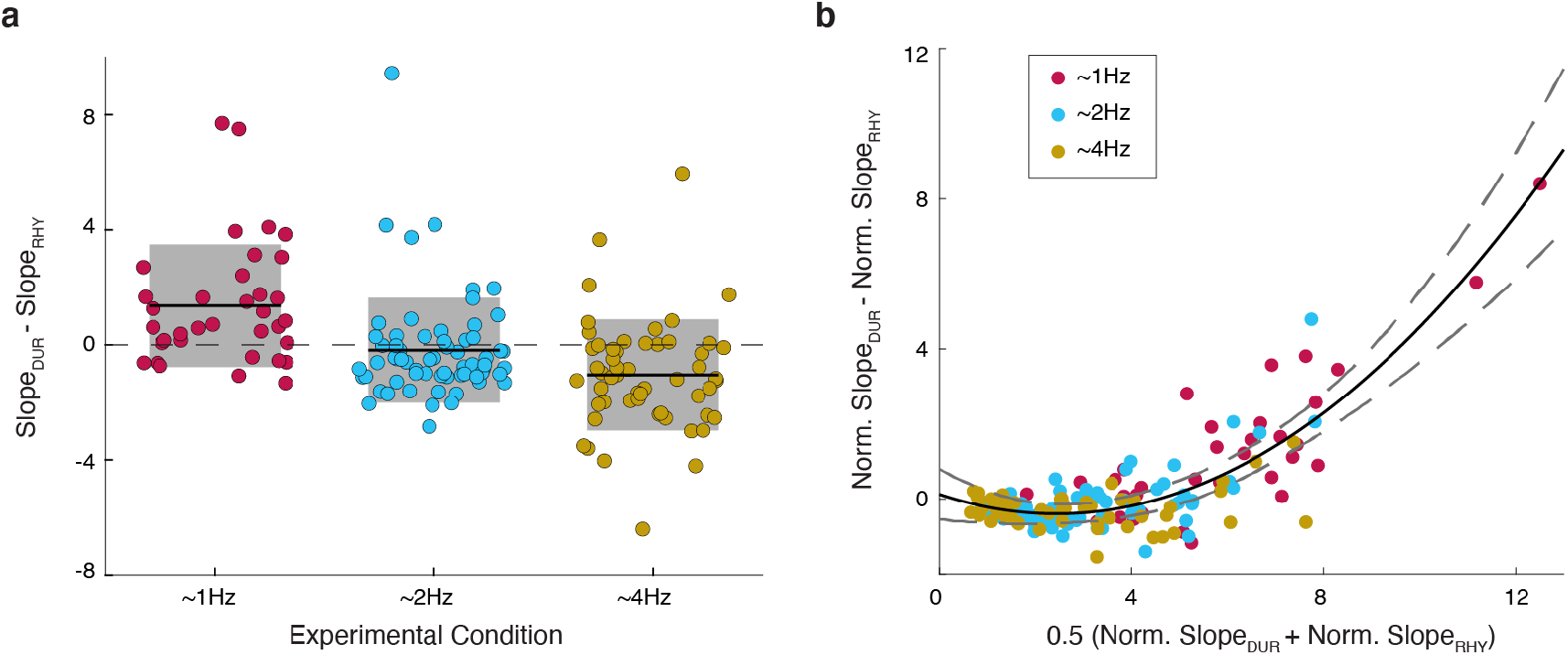
Behavioral responses. **a.** Algorithm preference for participants in the different experimental conditions. Difference of the slopes obtained by fitting a logistic regression for the participants answers as a function of the probe time computed from: the expected time according to the Duration (Slope_DUR_), or the expected time according to the Rhythm (Slope_RHY_) algorithms. Black line indicates the mean value and shaded region the standard deviation. **b.** Algorithm prevalence as a function of the overall performance. Slope difference between the two normalized logistic regressions as a function of the mean slope across algorithms. The black line represents a second degree polynomial fit and the dashed gray line the 95% Prediction Interval. The parameters of the fitted polynomial are: *y* = *ax*^2^ + *bx* + *c* with *a=0.087, b=-0.42, c=0.12*.

Next, we sought to understand if this effect is truly the result of a shift in frequency or is instead mediated by a third variable. Given that the attrition rate (participants removed because of noisy pattern of responses, see Methods) varied across studies (1.2Hz: 3%, 2Hz: 30%, 4Hz: 28%), with lower attrition for the condition preferring Duration algorithm, we considered it possible that the shift in algorithms was related to the shift in performance on the task across stimulus rates. Figure 3b confirms this hypothesis. We plot each participant’s algorithm preference (i.e., the difference in slopes) against overall performance (i.e. the sum of slopes) and found that the Duration method is preferred when performance is high. Furthermore, we found that the relationship between algorithms accuracy difference and overall performance can be modeled by a second degree polynomial (see Fig. 3b; adj. R-square = 0.71). The fitted polynomial dips below zero, confirming the Rhythm algorithm dominance for low performance.

### Bayesian Model

We sought to understand how these results would arise under optimal conditions, by creating a Bayesian model to simulate the temporal perception of the tones and decision making regarding whether the last tone was early or late. We built a model which treats the temporal perception of tones as a gaussian model. Each measurement records the tone’s timing with gaussian noise (See Figure 4a). The measurement is combined with a prior, representing the expectation of where the upcoming tone should be given the previous sequence. The prior is extrapolated from distributions of previous tones, whose timings and uncertainty are fed to an algorithm to predict the location of the next tone (Figure 4a, left). The algorithm can be used to make expectations either by Duration method or the Rhythm method, yielding different behavioral results in each case (Figure 4bc). The prior and measurement are combined in a multiplicative fashion to generate the posterior estimate of the true final tone. In the case of the final tone, the prior and likelihood are compared, yielding an estimate of how likely it is that the final tone should be considered early or late. The model’s true response is a weighted coin flip based on this probability.

**Figure 4.**
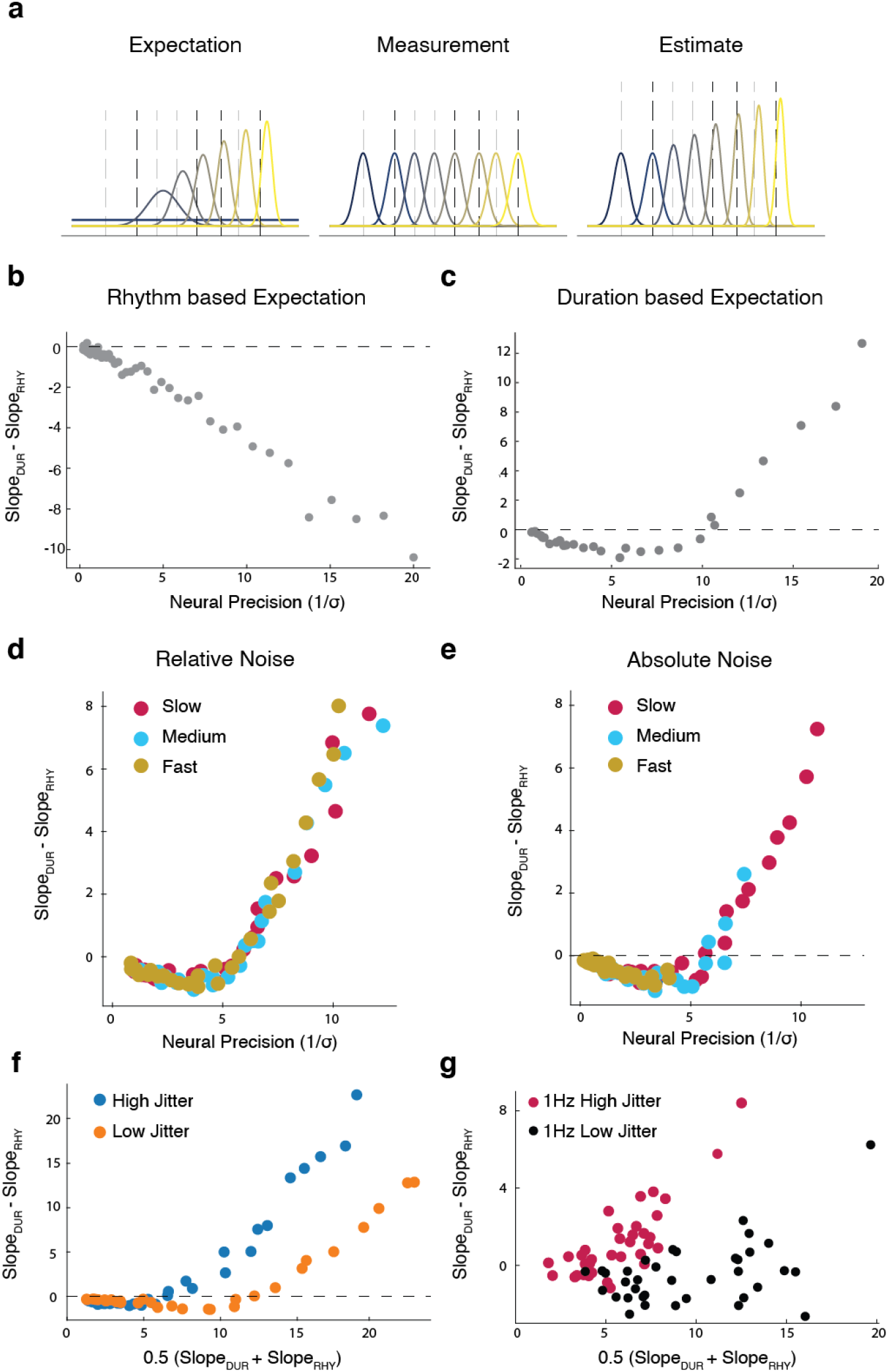
Bayesian model yields human-like performance. **a.** Example trial for temporal estimation of a series of tones in a single trial. The prior (left), likelihood (middle) and posterior (right) of each tone in the sequence. Distributions are color-coded from first to last tone in the sequence from blue to yellow. **b & c.** The differences in slope for the model when the prior is estimated using the Rhythm algorithm (**b**) or the Duration algorithm (**c**). **d & e**. Performance of the model for different stimulus rates corresponding to our human experiments where stimulus noise is relative to the stimulus rate (**d**) or the same across rates (**e**). **f.** Performance of the model when the temporal jitter in the tone sequences is altered. **g.** Performance of human subjects with lower temporal jitter. Red shows the original 1 Hz condition participants, black shows new participants with reduced jitter.

We change only the noise in the sensory measurement to mimic different participants in the experiment. In so doing, we find that the Bayes’ model with a duration-based prior yields remarkable similarity to the human behavior we found previously, showing both a preference for the rhythm algorithm under noisy conditions and a preference for the duration algorithm in clean ones. The rhythm-based prior yields a very different behavior, always preferring the rhythm algorithm. These findings lead us to conclude that participants are most likely using a duration-based estimate of their expectations and that the preference rhythm in low performance represents evidence of a prior adjusting timing toward rhythmicity when sensory measurements are noisy.

We next sought to understand how this model would explain the differences that we found across frequencies. We compared two versions of the model: one in which the sensory noise in the model was matched in an absolute sense (Figure 4e) across the stimulus rates, and another in which the noise was matched relative to the mean stimulus rate (Figure 4d), as would be predicted by Weber’s Law (CITE). The results suggest that differences only arise between algorithms when the sensory noise is considered to be absolute and not relative to frequency.

Lastly, we sought to test whether the model made any further predictions that could be tested by behavioral data. We found that if one reduces the jitter in time between participants, the model’s “Rhythm regime” expands (Figure 4f). We ran a fourth experiment to test this prediction, using the 1.2 Hz range but reducing the jitter in absolute terms to the values used in the 4 Hz experiment. We find that the data confirms the prediction of the model (Figure 4g). Despite better overall performance, the low-jitter experiment yields less Duration preference overall. We believe therefore that this model yields a straightforward understanding for how temporal prediction occurs under these scenarios.

### Adaptive Frequency Model

Having found a clear computational model for the behavior of our participants, we next sought to understand how neural dynamics could underlie such a model. We began by testing how well the phase of a neural oscillator, as typically described in the literature (see Methods for more detail), predicts early or late responses defined by Rhythm and Duration algorithms (see Figure S1a). We found that when the classic oscillator was driven by our ambiguous stimuli its phase significantly predicted the position of the last tone according to both algorithms, as long as the natural frequency of the oscillator matched the mean inter-tone interval of the stimulus. In addition, under these specific conditions (i.e., internal frequency congruent to the stimulus’ mean inter-tone interval) we found that manipulating the coupling parameter between the oscillator and the stimulus (see Figure S1a) modulated the model to better predict early or late according to one or another algorithm (see Figure S1b). Furthermore, by varying the coupling parameter the experimentally found relationship between overall performance and algorithm preference (i.e., better performance leading to Duration-algorithm preference, worse performance to Rhythm) was recovered (see Figure S1d). Despite these promising results, we found that when the natural frequency of the oscillator did not perfectly match the stimulus’s mean inter-tone interval, its phase prediction power decreased and the relationship between overall performance and algorithm preference vanished (see Figure S1d and e). While the oscillatory model clearly lacks key features needed to mimic human behavior (it fails to adjust to new stimulus rates), under very restricted regimes it contains the architecture required by human performance, namely to flexibly move between regimes more similar to rhythm and duration algorithms.

Based on this first result, we designed a structure for the oscillator model capable of adapting its natural frequency according to the temporal regularities of the stimulus. One of the parameters of the model represents the input coming from other brain regions. Crucially, the value of this parameter modulated the natural frequency of the system. Given that it is reasonable to assume a dynamic interaction between brain regions, we adopted a model in which the input that the oscillator receives from distant brain regions is a function of the perceived auditory period. Using this we were able to adjust the natural frequency of the oscillator on each new tone to match the average period of the current trial (Figure 5a). Furthermore, we allowed the model to learn across trials allowing it to stay in the space of probable stimulus rates given previous experience.

**Figure 5.**
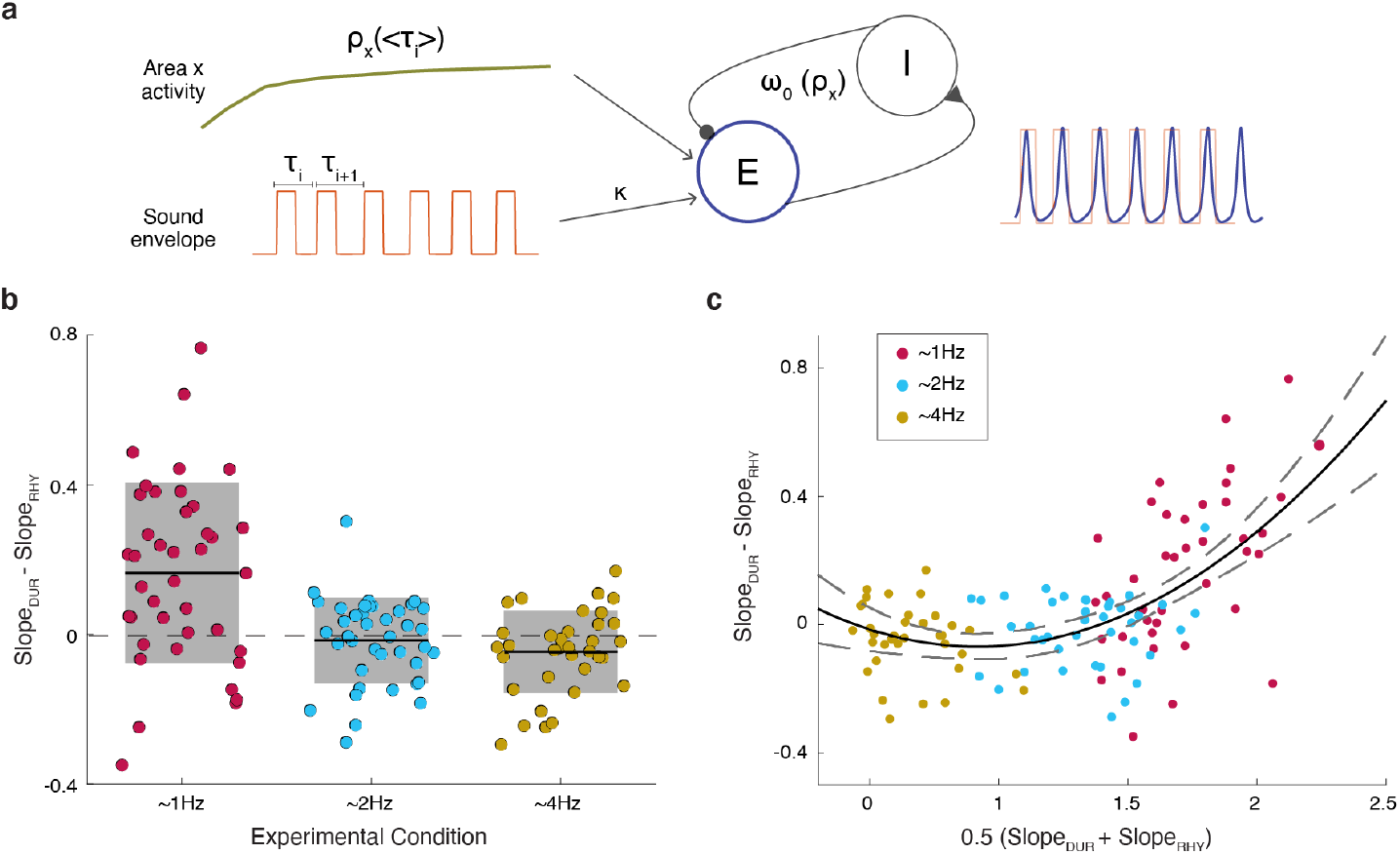
Adaptive Frequency Oscillator. **a.** Schematic of the AFO architecture. Populations of interacting excitatory (E, blue) and inhibitory (I) neurons constitute the oscillator (see Methods). The sound envelope (orange) drives the excitatory unit by some coupling gain (κ). The natural frequency of the oscillator is governed by input from an arbitrary area x (green) which stimulates E yielding faster or slower oscillatory input to match the mean period of the envelope. **b.** Algorithm preference for participants simulated with the AFO in the different experimental conditions. Difference of the slopes obtained by fitting a logistic regression for the answers computed according to the Duration (Slope_DUR_), or to the Rhythm (Slope_RHY_) algorithms as a function of the phase of the oscillator at the time of the probe (see Methods). Black line indicates the mean value and shaded region the standard deviation. **b.** Algorithm prevalence as a function of the overall performance. Slope difference between the two logistic regressions as a function of the mean slope across algorithms. The black line represents a second degree polynomial fit and the dashed gray line the 95% Prediction Interval. The parameters of the fitted polynomial are: *y* = *ax*^2^ + *bx* + *c* with *a=0.17, b=-0.26, c=0.05*. Simulated data mimic human behavior both in terms of preference for rhythm method in worse performance and matching the trend of stimulus rates.

With this design, we found that the *Adaptive Frequency Oscillator* (AFO) well mimics human behavior. The model yields both significant effects of frequency, shifting preference for duration and rhythm algorithms in a similar fashion to human performance (Figure 5b). At 1.2 Hz the Duration algorithm is preferred (Wilcoxon signed-rank test, two-sided p < 0.001), no significant difference between algorithms performance was found for 2 Hz (Wilcoxon signed-rank test, two-sided p =0.78) and at 4 Hz the Rhythm algorithm prevails (Wilcoxon signed-rank test, two-sided p =0.017). Also, the AFO yields a similar relationship between algorithm preference and overall performance (Figure 5c) as the experimental dataset. Oscillator’s phase better predicts early or late as computed by the Rhythm algorithm when overall performance is poor, while for enhanced performance it aligns better to the Duration algorithm estimations.

We sought also to compare this performance, at least qualitatively, to the behavior of a non-oscillatory temporal prediction. For this, we considered a state of the art temporal prediction model used to consider very short sequences, the Sensory Anticipation Module (SAM) from Eggers and colleagues^28^. SAM is a predictive ramping model that changes its speed so that the ramping value arrives at a threshold at the predicted expected moment. The module performs very well on the task and does match some of the behavior of human performance: specifically, it shows the switch in algorithm preference for the 4 Hz stimulus rate (**Figure S2**). However, this relationship between algorithm preference and stimulus rate is not monotonic as it was in human performance. Furthermore, algorithm preference is not related to overall performance as it was in human behavior and the bayesian model. Instead it is restricted to different frequency groups, yielding a different kind of behavior than what we found in our participants’ responses.

**Figure S1.**
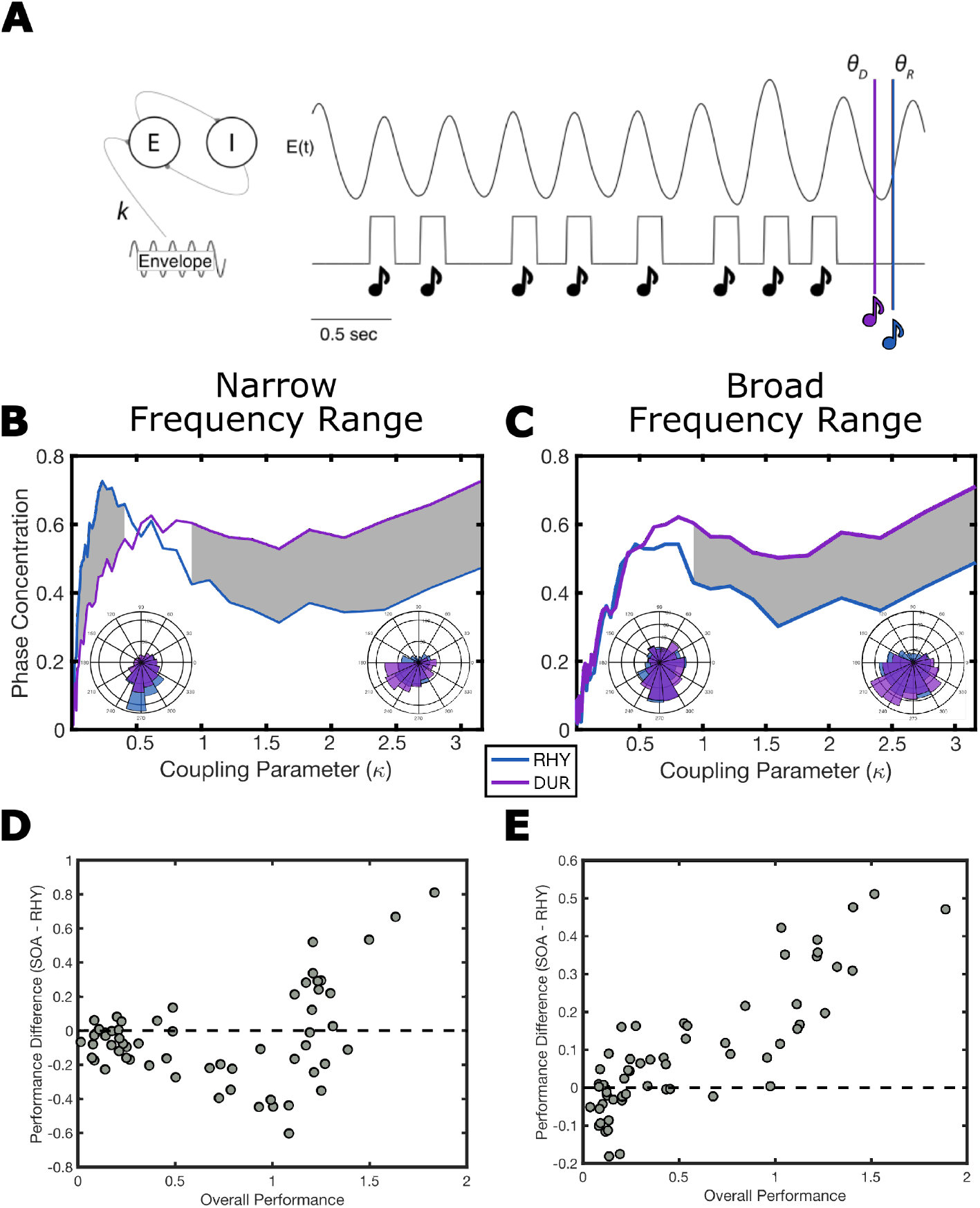
Classic oscillator performance. **a.** Set up of a Wilson-Cowan oscillator model (see Methods) with parameters set at: *a* = *b* = *c* = 10, *d* =− 2 and (ρ_*E*_, p_*I*_) = (1. 6, – 2. 9). The acoustic envelope of a stimulus trial drives the excitatory population with coupling determined by parameter *k*. The phase of the oscillator at the predicted time of the Duration algorithm (θ_*D*_) or the Rhythm algorithm (θ_*R*_) is extracted on each trial. **b.** Phase concentration of predicted phases, θ_*D*_ in purple and θ_*R*_ in blue across trials at a restricted range of stimulus rates (240 to 260 ms). Better phase concentration would lead to a more accurate prediction of the probe time relative to the corresponding algorithm. Shaded areas mark significant differences using the circular K test to test for significant differences in concentration (correcting for multiple comparisons using the false discovery rate Benjamini & Hochberg, 1995). Insets represent example concentrations at κ = 0.15, left, and κ = 2. 0. **c.** Same as **b** but with a range of stimulus rates that reflects the statistics of the experiment (210 to 290 ms). **d.** Algorithm preference of the Wilson-Cowan in the restricted range of stimulus rates (240 to 260 ms). The difference in slope parameters fitting a Logistic regression between phase of the oscillator at probe time and the correct response defined either by Duration or Rhythm algorithm (same as for the AFO model, see Methods). **e.** Same as D with the broader range of stimulus rates (210 to 290 ms).

**Figure S2.**
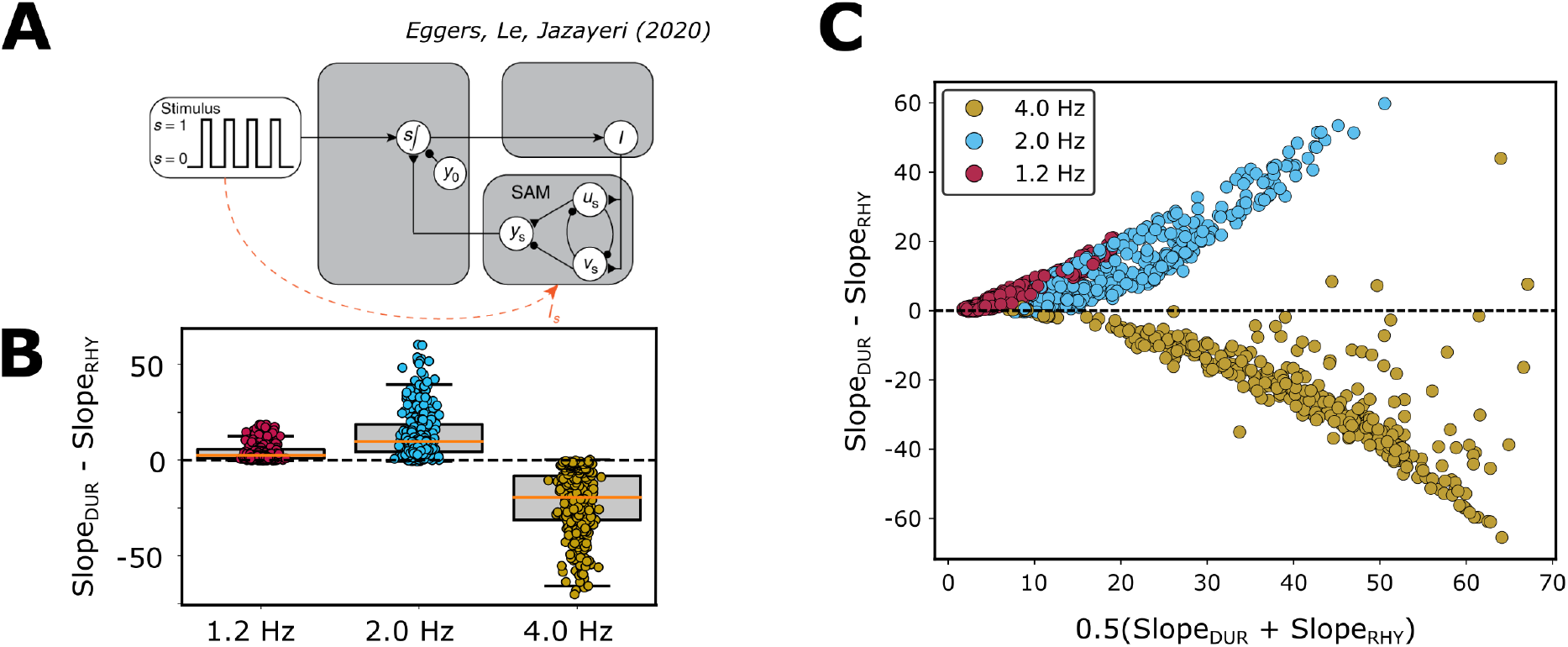
Performance of the Sensory Anticipation Module. **a.** Model schematic of the ramping model (adapted from Egger and colleagues^28^). The model contains two competing units *u_s_* and *v_s_* whose values decay to a stable point, driven by current *I* controlling the speed of this decay. Their difference yields *y_s_* a ramping value which is compared with threshold *y*_0_ at the time of a stimulus. The difference *d* = *y_s_* – *y*_0_ at the time of each tone is used to adjust *I* controlling the speed of the ramp to reduce *d* on the next interval. In our case, *d* is also used at the time of the probe tone to output a response to the behavioral trial. If *d* > 0, the model responds “late”; if *d* < 0, the model responds “early”. For code and further description of the module, see Eggers et al, 2020. **b.** The model responses are then treated as behavioral data. Late and early responses are coded as 1 and 0 respectively and a logistic function is fit and the slope extracted to identify the precision of the responses relative to the rhythm and duration algorithms. Slope differences are shown here by stimulus rates at 1.2 Hz (red), 2 Hz (blue) and 4 Hz (gold). **c.** Slope difference relative to overall performance for the same stimulus rates.

## Discussion

Our experiment uses a simple task design to study the added complexity of temporal prediction without perfect rhythmicity. We found that participants shift their method of expectation from rhythm-based to duration-based, with increasing overall performance. We then demonstrated that this behavioral pattern can be replicated by a Bayesian model of temporal perception, which yields further predictions also met by later experiments. Finally, we built an adaptive frequency oscillator that adjusts its spontaneous rate to match the overall period of the trial. We found that this model mimicked all of the behavior of our human participants, moreso than a state-of-the-art predictive ramping model. Our findings push proponents of oscillator models in perception to move beyond basic oscillators to deal with complex stimuli, acknowledging that such units must be placed within the larger neural system. Crucially, at the same time, we validate the use of oscillator-based models as a potential means of temporal inference, considering the phase adjustments in response to input as a kind of neural heuristic for a prior expectation for rhythmicity in a Bayesian sense.

### Absolute vs Relative Timing

We have devised the Duration and Rhythm algorithms of temporal estimation to relate directly to Absolute and Relative Timing mechanisms discussed in the literature. However, It is important to note that these algorithms are not the only possible ones. Nor do we claim that a participant who is doing better with one algorithm definitively relies on it to perform temporal estimation. Instead, we consider these two methods as goal posts, ways to measure any possible algorithm as more similar to one or the other.

Previous work has considered the two timing mechanisms, absolute and relative, as being supported by two connected timing systems, such that absolute timing is primarily used on irregular sequences and relative timing used on regular/rhythmic sequences. Teki and colleagues^24^ developed a unifying theory in which mechanisms associated with relative timing yield a rough temporal estimate based on recently heard intervals and those associated with absolute timing yield error correcting measurements to refine the measurement. By this theory, as irregular sequences have greater error, absolute timing is more heavily relied upon, whereas the reverse is true for regular sequences. While this theory has proved fruitful, we believe it can be refined by considering how the relative timing mechanism yields predictive estimates during irregular sequences. Our modeling results suggest that the two mechanisms might instead be combined not as an error correction process but as the combination of prior and sensory measurement to yield a final temporal estimate. In this sense, the relative timing mechanism may fit as a prior expectation and the absolute mechanism as a likelihood or raw sensory measurement, both estimations combined in a Bayesian sense to yield a final estimation.

Furthermore, from a dynamical perspective, it may be tempting to consider the rhythm algorithm to relate directly to oscillations and the duration method to non-oscillatory mechanisms (e.g. ramping). Our results refute this possibility. We find both scenarios in which oscillators yield either rhythm or duration like behavior and ones in which ramping mechanisms do the same. As such, we caution the reader from making this kind of overly simplistic link between human behavior and possible neural mechanisms. In fact, we find that an adaptive frequency oscillator is capable of flexibly navigating the two timing mechanisms representing a unifying theory.

### Human Behavior

Our analysis showed that participant behavior varies between these two anchor algorithms. This variation was explained by two major features: the stimulus rate (slower rates were more likely to yield performance similar to the duration algorithm) and overall performance (participants’ better overall performance was associated with the duration algorithm). Furthermore, overall performance seems to be the dominating feature explaining algorithm preference, since all stimulus rates can be readily described in terms of the relationship between performance and model preference using a single parabolic fit. The effect of stimulus rate may be best considered as placing the population on a different part of the function between performance and model preference.

This single fit across stimulus rates is found in the context of the well-known Weber’s law in which overall performance is considered relative to the duration of intervals being perceived. We opted to consider our analysis in this context as is commonly done. Therefore, in this case, the jitter of the probe tone is normalized by the mean tone duration of the experiment. That normalization yields such alignment across stimulus rates confirms its validity as a means of comparing across frequencies.

### Bayesian Modeling

To better understand human performance, we developed a Bayesian computational model to see how optimal estimation would occur. We related participants’ behavior to prior expectations and inferences in a Bayesian model and predicted that they should match if the internal model is statistically optimal. Our findings suggest that participant behavior is consistent with Bayesian estimation assuming Duration-based (and not Rhythm-based) priors. It therefore explains the effect of significant preference for the Rhythm method at worse performance levels as increasing dominance of the prior in the cue tone phase under scenarios with high sensory noise. When participants are more uncertain about their own sensory measurements, the expectation of similarity between intervals yields a stronger effect of rhythmicity in the estimation.

Interestingly, the Bayesian model makes an added prediction that we had not initially anticipated. As precision in the tone sequence increases, making the sequence closer to isochronous, the function that governs the relationship between overall performance and model preference flattens, expanding the range in which the rhythm model performs best. We ran a fourth experiment using the stimulus rates from the 1.2 Hz experiment but the jitter of the 4 Hz experiment (in absolute terms). Comparing this experiment with the initial 1.2 Hz experiment confirmed the predictions of the model. Showing a reduction in duration preference despite higher overall performance. We feel therefore that this model well reflects human behavior and further validates our behavioral findings. It also confirms previous predictions suggested by Teki and colleagues^24^ that more regular sequences rely more heavily on relative timing mechanisms.

### Implementational models

We found that an adaptive frequency oscillator model well reflected the participants’ behavior and the Bayesian model predictions. This model, in essence, combines elements of other models that often have been used for temporal prediction analysis, in particular, oscillator models and predictive ramp models. In a supplementary analysis, we found that a simple oscillator could not on its own replicate human behavior in response to ambiguous rhythms, unless the mean rate of the input was near the spontaneous rate of the model. In this restricted scenario, the oscillator could successfully switch between rhythm and duration models depending on its coupling strength, and subsequently overall performance.

Meanwhile, the predictive ramp model that we tested, the Sensory Anticipation Module (SAM) designed for use in a tapping study, is able to successfully make reasonable predictions at a wide array of stimulus rates. Furthermore, it showed differences in model preference based on stimulus rate. However, overall performance seemed to have no bearing on the model preference, unlike human behavior. Therefore, while the architecture of SAM is clearly well designed and reflects many aspects of human behavior, it appears to miss something in order to truly reflect human performance.

Our AFO model is meant as a coarse amalgamation of these two models. Using the strengths of each model to ameliorate the weaknesses of the other. AFO contains the oscillatory component which allows for shifting in model preference on the basis of performance, and also the adjustment of speed (or period in this case) provided by the architecture of SAM. The combination of the two yields simulations that well resemble human behavior both in terms of behavior across stimulus rates and overall performance, making it a viable candidate for a neural mechanism to underlie temporal prediction.

One potential advantage of the AFO is that it also allows for input to adjust frequency not only based on the current stimulus but more abstract information. Added excitatory or inhibitory input from top-down/frontal inputs could theoretically adjust the expected frequency based on prior knowledge: expectation that the sequence will speed up based on previous tendencies, or the recognition of more complex hierarchical musical rhythms. Such input would fit well with recent work by Cannon and Patel^29^ which proposes a looping pattern in Supplemental Motor Area (SMA) whose speed is governed by input current. While their proposal is more specific regarding anatomy, we feel that this concept fits well with the AFO framework. Similarly, Rimmele and colleagues^30^ proposed oscillatory dynamics in the auditory cortex as a local hard-coded constraint that could be manipulated by top-down control via phase-reset (rather than frequency shift) most likely from cortical motor areas, cerebellum, or basal ganglia.

In addition, another advantage of AFO over the classical oscillator is the reduction of phase precession. When the classical oscillator model is driven by a periodic input, even when it synchronizes, the phase at which the locking takes place is modulated by the stimulus’ rate relative to the oscillator’s natural frequency^31^. This kind of phase shift is potentially problematic for a model that uses phase as an index of temporal prediction. The AFO postulated in this work overcomes this issue. Given that the natural frequency of the system is adjusted to match the external one, the oscillator synchronizes at zero phase lag for every external rate. Still, AFO does not represent a full account of the neural mechanism, taking several shortcuts to ease computational load and complexity. For example, the model adjusts its own period to match the mean interval of the period without explaining how this mean would be estimated. Future work will determine how this period matching is assessed and in what neural anatomical regions.

Still, our study shows that an oscillator component within a more complex model explains aspects of human behavior that have not yet been explained without it. Such work is in line with advancements in adjacent fields developing the notion of oscillatory behavior into more complex/realistic networks with great success^13,32–34^. Of these, particularly noteworthy are models of perception as phase locked loops (PLL)^32,35–38^, which use top-down feedback to adapt sensory organs and maintain a fixed phase relationship between input and neural activity. In our case, rather than overtly adapt the oscillator’s phase, we focused on matching the oscillator’s period assuming that synchronization dynamics will yield meaningful phase computations. This choice derives from the fact that our stimuli are discrete units without the continuously varying dynamics of other natural stimuli such as speech for which PLL was initially designed.

### Relating oscillators and Bayes’

How the AFO model successfully mimics Bayesian inference remains unclear. However, an implementational model of Bayes requires the ability to combine expectation and sensory estimate weighted by the uncertainty of each. The oscillator component of AFO clearly contains these features. The phase of the oscillator represents an expectation, the sound envelope represents sensory estimate, and the coupling strength controls the weighting of each to yield a phase reset of appropriate magnitude. When coupling strength is low, sensory measurements are given low weight and the pre-stimulus phase/expectation is dominant. When coupling strength is high, oscillator phase is heavily influenced by the stimulus input yielding strong phase resets that may not be phase dependent. Whether this coupling strength is expressly manipulated as a result of some meta knowledge regarding the system’s own uncertainty, or is instead a bottom-up feature in which noisier measurements influence phase adjustments less remains an open question.

We can not definitively say whether oscillators in some form are necessary at the implementational level to yield this Bayesian behavior in regular sequences. Still, it is reasonable to think an oscillator may be one of the most plausible options. Recent work from Sohn and colleagues^16^ showed that Bayesian estimates of single intervals can be represented in neural populations by a curvature in the population’s state-space trajectory. Such curvature when placed in a sequence of events would likely become a repeating loop, virtually indistinguishable from the limit cycle that would represent oscillatory behavior. In the expectation of regular sequences, oscillator behavior may be advantageous.

This work also raises the interesting – and, to our knowledge, overlooked – prospect that certain neural oscillators might be considered as *hard-coded* priors to subserve temporal predictions. Forward models are generally thought of as being built and optimized on the basis of an individual’s prior sensory experience, through learning. Although acquired sensory experience clearly matters to build such models, the current work allows speculation that innate mechanisms, although not of a top-down nature per se, might contribute to the inferential process. It may be that some neural oscillations reflect the biological internalization of environmental statistics within the neurosensory apparatus. Whether intrinsic neurophysiological mechanisms are inherited or whether they gradually adapt to the statistics of the environment through learning remains an open question.

### Conclusion

Taken together, our findings show that participants process imprecise rhythms in a manner consistent with Bayesian principles and primarily use a Duration estimate method (expecting drift) to build their expectations. Remarkably the preference for the Duration algorithm (with better performance) occurs even though the stimuli themselves were generated by the Rhythm algorithm. Adding to this computational account for the behavioral pattern, we found that an adaptive frequency oscillator represented a candidate neural mechanism which well replicated human performance. This model represents a significant advance which we think should guide the development of future oscillatory models of temporal prediction. In this case, the frequency adaptation represents a mechanism for estimating the period, while the synchronization reflects successful estimation of timing in this period as a kind of neural heuristic for a Bayesian prior. Furthermore, it suggests that frequency adaptation may be a key optimization to the oscillator hypothesis for sensory prediction. Altogether, this work advances a convergence between oscillatory and Bayesian models of perception in time, in which oscillatory components behave as a rhythmic prior, using phase adjustment as a neural heuristic for prior updates to sensory measurements.

## Code availability

All computer code used for this study is available upon request.

## Data availability

All data needed to evaluate the conclusions are present in the paper. Additional data related to this paper may be requested from the authors.

## Acknowledgment

The authors would like to thank David Poeppel for his comments and helpful discussion. This work was supported by UNAM-DGAPA-PAPIIT IA202921 (M.F.A), IBRO Return Home Fellowship (M.F.A.), Fondation Fyssen Postdoctoral Fellowship (K.B.D.)

## Author contributions

K.B.D. & M.F.A. conceived the project, ran simulations, analyzed and collected the data. L.H.A. and M.F.A. supervised this work. All authors wrote the manuscript.

